# Distinct roles of the *Lyve1* lineage in heart development

**DOI:** 10.64898/2026.01.06.697888

**Authors:** Konstantinos Klaourakis, Karolina Zvonickova, Jacinta Kalisch-Smith, Nicola Smart, Duncan Sparrow, David G. Jackson, Paul R. Riley, Joaquim M. Vieira

## Abstract

*Lyve1^Cre^* is widely used to study lymphatic endothelial cells, but its activity in the embryonic heart has not been comprehensively defined. Here we show that the *Lyve1* lineage contributes to multiple cardiovascular cell types, including venous endothelium of the sinus venosus forming coronary vessels, endocardial cells generating complementary coronary territories and valve structures, lymphatic endothelial cells required for cardiac lymphatic development, and tissue-resident macrophages within the embryonic myocardium. Functional perturbations demonstrated that *Notch1* gain-of-function in *Lyve1^+^* lineages disrupted vascular development, while *Prox1* deletion caused severe edema, blood-filled lymphatics, and perinatal lethality. Mid-gestation mutants exhibited underdeveloped, blood-filled lymphatics despite intact blood vasculature, and by late gestation cardiac lymphatics were depleted. These findings reveal distinct cardiovascular roles of the *Lyve1* lineage and establish *Lyve1^Cre^* as a versatile but non-specific tool, underscoring the need for careful interpretation when investigating heart development and regenerative processes.

## Introduction

The lymphatic vasculature plays a fundamental role in vertebrate physiology, maintaining tissue fluid balance, enabling immune cell trafficking, and supporting lipid absorption^1,2^. Its developmental origins have, therefore, been a central focus of vascular biology, and in particular regarding the specification and diversification of lymphatic endothelial cells (LECs)^1,3–6^. In mouse, LECs are specified during embryogenesis from paraxial mesoderm–derived ETV2^+^ angioblasts and endothelial cells of the anterior cardinal vein, beginning around embryonic day (E) 9.5^4,6,7^. These precursors, defined by expression of the transcription factor PROX1, generate the majority of the lymphatic endothelium, including cardiac lymphatics that emerge after E12.5^5,8^. Additional progenitor sources, such as hemogenic endothelium, the second heart field (SHF), and dermomyotome have also been implicated in contributing to the LEC pool^4,8–10^. Expression of *Prox1* is both necessary and sufficient for LEC specification, and conditional deletion of *Prox1* has been widely used to interrogate lymphatic development, as constitutive *Prox1* knockout mice fail to survive beyond E14.5^8,9,11^.

Loss-of-function studies employing transgenic Cre drivers (*Tie2-Cre*, *Vav1-Cre*, *Isl1-Cre*, *Pax3-Cre*) have demonstrated that venous-, hemogenic-, SHF- and paraxial mesoderm-derived LECs are each required for cardiac lymphatic development^4,8–10^. However, transgenic Cre strategies are limited by integration site effects, copy number variation, and epigenetic alterations that can alter Cre expression patterns^12,13^. Knock-in Cre lines, in which recombinase activity is driven by endogenous regulatory elements, provide a more faithful alternative and have become widely adopted in developmental biology^12,13^.

*Lyve1^Cre^* is a knock-in line generated under the control of the endogenous *Lyve1* promoter and is extensively used as a genetic tool to study lymphatic biology^14–22^. Reported activity includes yolk sac hemogenic endothelium, embryo-derived hematopoietic precursors, and subsets of blood endothelial cells, in addition to LECs^14–16^. Despite its broad application, *Lyve1^Cre^* activity in the embryonic heart has not been comprehensively defined. This uncertainty is particularly relevant given the emerging importance of cardiac lymphatics in myocardial homeostasis, repair, and regeneration^8,23–27^. Recent studies have highlighted the role of lymphatic vessels in clearing inflammatory mediators, supporting immune resolution, and facilitating tissue repair after injury, underscoring the translational potential of understanding their developmental origins^26,27^.

A further consideration is that Cre drivers are often used not only for lineage tracing but also for conditional gene deletion. In this context, the specificity of *Lyve1^Cre^* becomes critical, as recombination in non-lymphatic lineages could confound interpretation of phenotypes. Thus, defining the full spectrum of *Lyve1* activity is essential for accurate use of this line in developmental and regenerative studies.

Here, we used *Lyve1^Cre^* to conditionally delete *Prox1* and assess its role in cardiac lymphatic development. Genetic lineage tracing revealed that *Lyve1^Cre^* marks multiple cardiovascular cell types, including venous endothelium of the sinus venosus, endocardial cells contributing to coronary vessels and valve structures, lymphatic endothelial cells required for cardiac lymphatic development, and tissue-resident macrophages populating the embryonic myocardium. Functional experiments demonstrated that *Notch1* gain-of-function in *Lyve1Cre^+^* cells disrupted vascular development, while conditional deletion of *Prox1* resulted in oedema, blood-filled lymphatics, and perinatal lethality. By E16.5, mutant hearts contained underdeveloped lymphatics filled with blood despite intact blood vasculature, and by E18.5 cardiac lymphatics were depleted entirely.

Together, these findings establish *Lyve1^Cre^* as a versatile but non-specific tool for lineage tracing and genetic manipulation in the heart. They highlight the distinct cardiovascular roles captured by *Lyve1* activity and emphasize the need for careful interpretation of phenotypes in developmental and regenerative contexts. Within the broader field of lymphatic vessel biology, our work provides insight into the intersection of lymphatic and cardiovascular development, with implications for understanding fundamental mechanisms of morphogenesis and for translational approaches to optimize cardiac repair.

## Results

### *Lyve1^Cre^* marks cardiac lymphatics and coronary vessels during embryogenesis

*Lyve1^Cre^* has previously been reported to label coronary blood endothelium, consistent with widespread LYVE1 expression in the embryonic vasculature^28^. To investigate the activity of the *Lyve1*^+^ lineage in the developing heart, *Lyve1^+/Cre^* mice were crossed with R26R-tdTomato reporters, and embryonic hearts were harvested at E14.5 and E16.5 (Figure 1). At E14.5, immunostaining of *Lyve1^+/Cre^;R26R-tdTomato* hearts with PECAM1, LYVE1, and PROX1 revealed Cre activity in both blood and lymphatic vasculature (Figure 1A–L). tdTomato signal co-localized with PECAM1 in large vessels and capillaries across the ventral surface (Figure 1D), and in PECAM1^+^ vessels of the sinus venosus (SV) and ventricular walls on the dorsal side (Figure 1G,J). Blood vessels lacked LYVE1 immunoreactivity at this stage, suggesting transient *Lyve1* expression earlier in development. On the ventral side, developing PROX1^+^ lymphatic vessels were either LYVE1^−^ near the outflow tract or LYVE1^+^ towards the apex (Figure 1B, E–F), with both populations expressing tdTomato. In contrast, PROX1^+^ LECs near the SV were negative for LYVE1 and tdTomato, with *Lyve1* labelling only evident in lymphatic vessels extending into the base of the heart (Figure 1H, K–L). By E16.5 (Figure 1M-T), staining with the venous marker EMCN confirmed widespread tdTomato expression in coronary veins and capillaries from the SV to the ventricular wall and apex (Figure 1M–O, R). At this stage, all cardiac LECs co-expressed LYVE1 and tdTomato (Figure 1M, P–Q, S–T).

**Figure 1.**
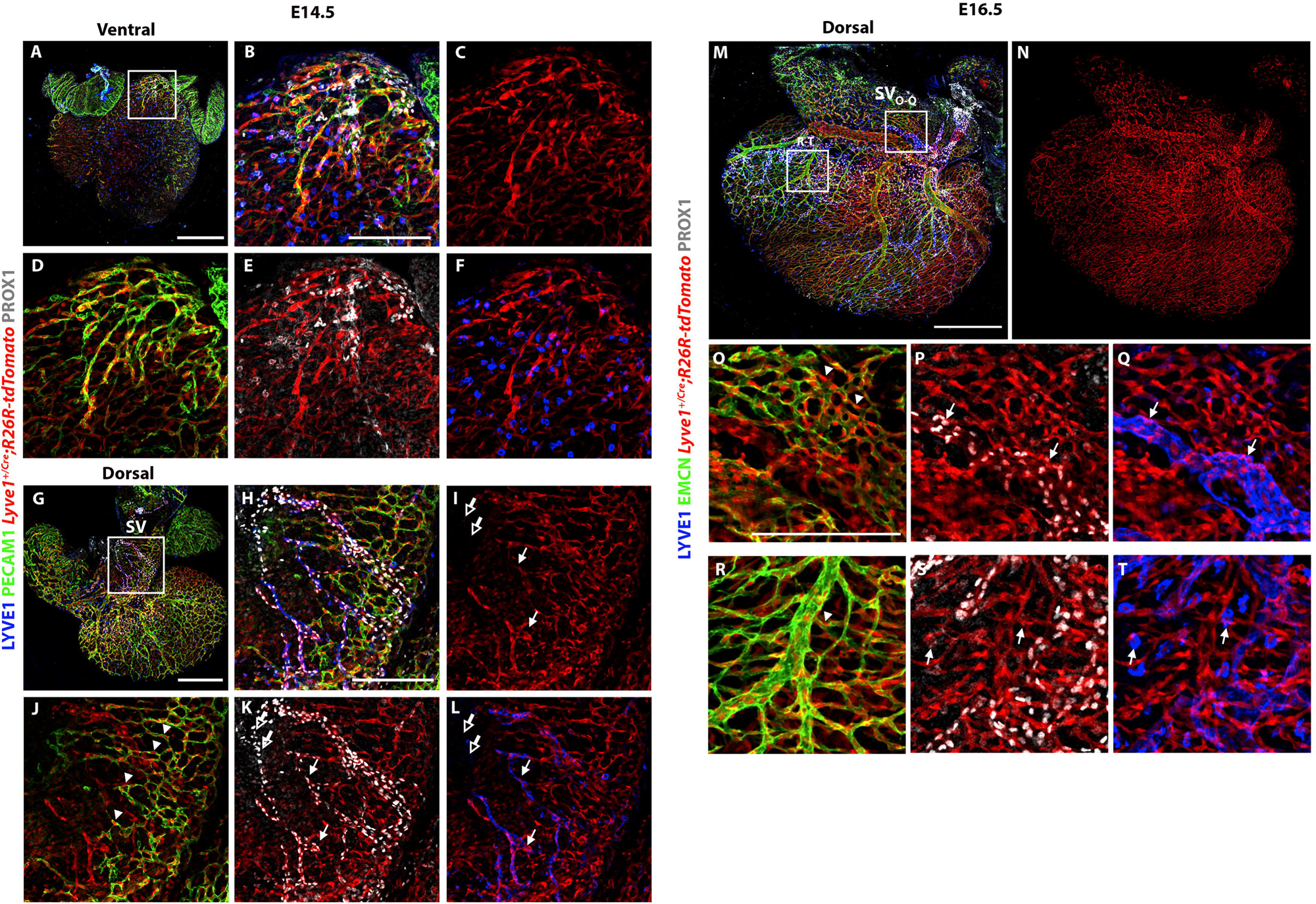
Confocal imaging of *Lyve1^+/Cre^;R26R-tdTomato* embryonic hearts reveals Cre activity in coronary vasculature and developing lymphatic vessels. Visualization of the ventral side of the heart at E14.5 (A-F). PECAM^−^;PROX1^+^;LYVE1^+^;tdTomato^+^ precursor LECs start forming lymphatic vessels (B-C and E-F). tdTomato is also visible in PECAM^+^;PROX1^−^;LYVE1^−^ coronary vessels (B-D). Visualization of the dorsal side of the heart at E14.5 (G-L). Near the SV, PROX1^+^ LECs appear to be negative for LYVE1 and tdTomato (hollow arrows), contrary to LECs near the tips of the vessels which are LYVE1 and tdTomato positive (H-I and K-L; white arrows). Coronary vessels in the SV are also positive for tdTomato (I-J; arrowheads). Visualization of the dorsal side of the heart at E16.5 (M-N, O-Q and R-T). At this timepoint all LECs appear to be marked by *Lyve1^Cre^*, both near the SV and towards the apex (white arrows). tdTomato signal is detectable in both large coronary veins and capillaries stained with EMCN, although they are LYVE1 negative (M-O, R; arrowheads). B-F magnified view of A box; H-L magnified view of G box; O-Q and R-T magnified views of M boxes. n = 5 for each time point. Scale bars: 0.5 mm for A,G and M-N ; 0.2 mm for B-F, H-L and O-T.

Overall, these results suggest that *Lyve1* is transiently expressed by the coronary vasculature before E14.5, and that not all cardiac LECs are marked by *Lyve1^Cre^*, particularly in the vicinity of the SV.

### *Lyve1^Cre^* labels the common cardinal vein and intersomitic vessels

During embryogenesis, coronary vessels arise from two progenitor populations that contribute to complementary regions of the heart^29–33^. Venous endothelium of the SV gives rise to vessels in the dorsal and lateral outer myocardial wall, while the endocardium contributes to the inner myocardial wall, interventricular septum, and ventral midline^29–33^. To assess *Lyve1^Cre^* activity in venous endothelium, *Lyve1^+/Cre^;R26R-tdTomato* embryos were stained for PROX1 and EMCN at E10.5 (Supplementary Figure 1A-M). tdTomato co-localized with EMCN in cardiac capillaries (Supplementary Figure 1A–E) and intersomitic vessels (ISVs; Supplementary Figure 1A,J–M). In the cardinal vein (CV) and ISVs, precursor LECs were either PROX1^+^;EMCN^+^;tdTomato^−^ or PROX1^+^;EMCN^+^; tdTomato^+^, indicating that *Lyve1^Cre^*does not label all cardiac precursor LECs or is activated later in specification (Figure 6.1B,H). The spatial distribution of CV and ISV precursor LECs supports active migration toward the SV and outflow tract to form the first cardiac lymphatics at E12.5^8^. PROX1 expression was also detected in the heart, consistent with prior reports^34^. Additionally, scattered PROX1^−^;EMCN^−^;tdTomato^+^ cells were observed throughout the embryo, likely yolk-sac–derived macrophages (Supplementary Figure 1A).

To further examine *Lyve1^+^* lineage contributions, *Lyve1^+/Cre^;R26R-tdTomato* embryos were harvested at E12.5, sectioned to reveal jugular veins (JVs) and jugular lymph sacs (JLSs), and stained for EMCN and PROX1 (Supplementary Figure 2). tdTomato+ cells were present in both JV endothelium (PROX1^−^;EMCN^+^) and JLS lymphatic endothelium (PROX1^+^;EMCN^−^), confirming *Lyve1Cre*-labelled contributions to blood and systemic lymphatic vasculature (Supplementary Figure 2A–M).

Taken together, these findings demonstrate that *Lyve1* is expressed during early embryogenesis in venous endothelium contributing to SV-derived coronary vessels, as well as systemically in the CV and developing ISVs, the sources of cardiac lymphatics.

### *Lyve1^Cre^* labels the endocardium during embryogenesis

Coronary vessels of the interventricular septum and ventral mid-portion of the heart arise from endocardial-derived sprouts^31–33^. To examine *Lyve1^Cre^* activity in the endocardium, *Lyve1^+/Cre^;R26R-tdTomato* hearts were harvested at E14.5 and E16.5, sectioned transversally, and stained for PECAM1 and PDPN (Figure 2A-L). At both stages, tdTomato was detected in the PDPN-expressing epicardial layer, either in macrophages (PECAM1^−^; Figure 2C) or endothelial cells (PECAM1^+^; Figure 2I). tdTomato expression was also observed in developing cardiac valves (Figure 2D,J) and the interventricular septum (Figure 2F,K). Notably, tdTomato+ cells in the valves were PECAM1^−^, consistent with previously described endocardium-derived tissue-resident macrophages essential for valve remodeling^35^. Finally, tdTomato was abundantly expressed in endocardial trabeculae alongside PECAM1 (Figure 2F,L), supporting *Lyve1^Cre^* activity in the developing endocardium and its contribution to endocardial-derived coronary vessels.

**Figure 2.**
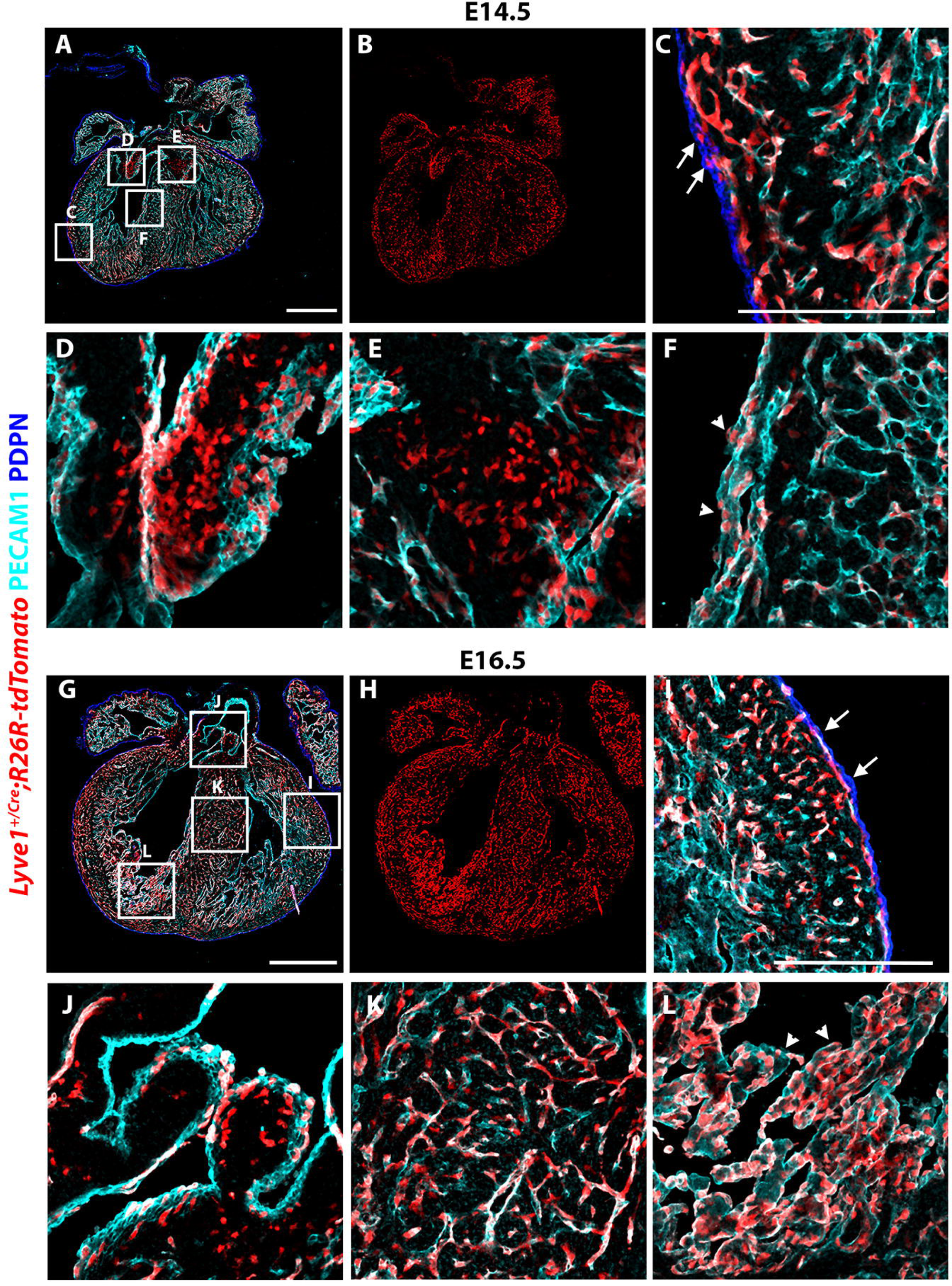
*Lyve1^Cre^*is activated in the developing endocardium and cardiac valves. Cryosection of *Lyve1^+/Cre^;R26R-tdTomato* heart at E14.5 shows expression of tdTomato throughout the embryonic heart (A-B). High magnification images from different areas of the E14.5 heart (C-F). tdTomato is detected in PECAM1 stained vessels in the myocardium, as well as in PECAM1 negative cells, possibly macrophages (white arrows), in the PDPN-expressing epicardium (C). Macrophages residing in cardiac valves appear to express tdTomato (D-E). The endocardial layer clearly co-expressed tdTomato and PECAM1 (F; arrowheads). Cryosection of *Lyve1^+/Cre^;R26R-tdTomato* heart at E16.5 (G-H). Vessels expressing tdTomato are detectable in the sub-epicardial layer at this timepoint (I; white arrows). Macrophages expressing tdTomato near the cardiac valves (J). In both the interventricular septum and the endocardium PECAM1 is found only in cell that co-express tdTomato, representing coronary vessels and trabeculae (K-L; arrowheads). C-F magnified views of A boxes; I-L magnified views of G boxes. n = 5 hearts per stage. Scale bars: 0.5 mm for A-B, G-H; 0.2 mm for C-F and I-L.

### *Lyve1^Cre^*-driven activation of *Notch1* disrupts embryonic vascular development

To test whether *Lyve1^Cre^* targets the developing endothelium and endocardium, *Notch1* was ectopically activated by crossing *Lyve1^+/Cre^* mice with *Gt(ROSA)26Sor^tm1(Notch1)Dam^*(N1ICD)^36^. The conditional N1ICD line expresses the Notch1 intracellular domain upon Cre activation. Previous studies using *Tie2-Cre;N1ICD* embryos reported severe vascular defects, including disorganized, low-density vessels in the head and disruption of cardiac morphogenesis, with lethality by E10.5^37,38^. Accordingly, *Lyve1^+/Cre^;N1ICD* embryos were examined at E10.5. Whole-mount EMCN staining and confocal imaging revealed abnormal vascular density, with thicker, disorganized vessels compared to controls (Supplementary Figure 3A–F). The phenotype was most pronounced in head capillaries and ISVs, resembling *Tie2-Cre;N1ICD* embryos at E9.5^37,38^. In contrast, no gross cardiac morphological defects were detected in *Lyve1^+/Cre^;N1ICD* mutants, which may reflect technical limitations of whole-mount imaging or differences in spatial and temporal *Cre* activity between *Tie2* and *Lyve1* drivers.

Overall, these findings demonstrate that *Lyve1^Cre^* activation of *Notch1* disrupts embryonic vascular development, supporting *Lyve1* expression in blood endothelium during early embryogenesis.

### *Lyve1^Cre^* marks tissue-resident cardiac macrophages during organogenesis

Macrophages first appear in the heart around E10.5, migrating from the yolk sac to the epicardium/subepicardium compartment^23,39^. LYVE1 is a known tissue-resident macrophage marker, but its activity in cardiac macrophages has not been systematically assessed. To address this, *Lyve1^+/Cre^;R26R-tdTomato* embryos were harvested at E14.5, and hearts stained for CD68 (pan-macrophage marker), CD206 (tissue-resident marker), and LYVE1, followed by confocal imaging (Figure 3A–L). tdTomato and CD206 were detected in nearly all CD68^+^ cardiac macrophages, with both LYVE1^+^ and LYVE1^−^ subsets present on the ventral and dorsal heart surfaces (Figure 3B–F, H–L). Quantification revealed that ∼54% of macrophages were CD68^+^;CD206^+^;tdTomato^+^;LYVE1^+^, while ∼44% were CD68^+^;CD206^+^;tdTomato^+^; LYVE1^−^. Overall, 98% of cardiac macrophages expressed tdTomato, but only 44% expressed LYVE1, suggesting *Lyve1* activity during early hematopoiesis (Figure 3M). A minor population (2%) of CD68^+^;CD206^−^;tdTomato^−^;LYVE1^−^ macrophages was also identified (Figure 3M), potentially representing fetal liver monocyte-derived macrophages^40^.

**Figure 3.**
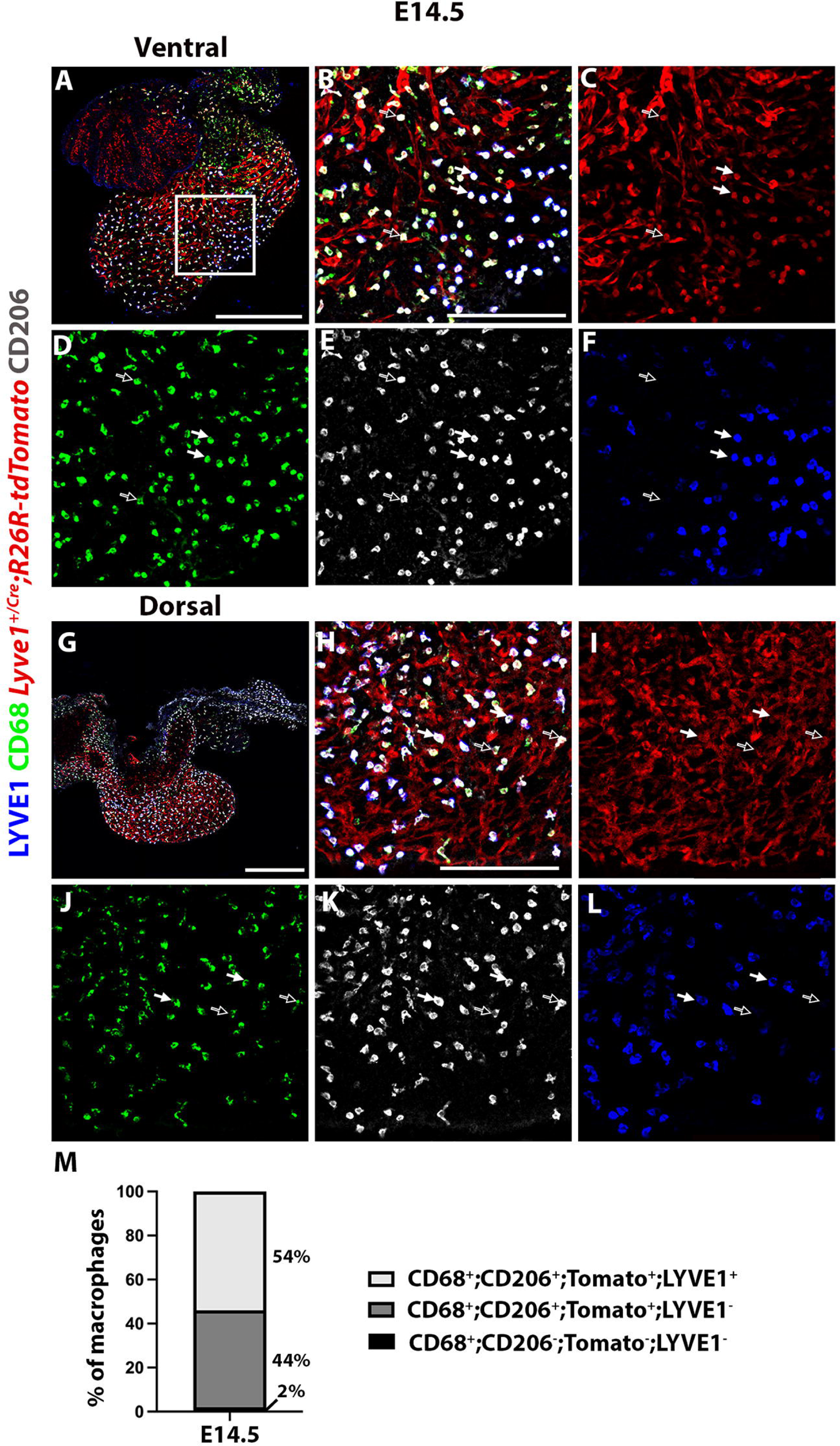
*Lyve1^Cre^* is activated in cardiac resident macrophages at E14.5. Macrophages visualized on the ventral aspect of the heart of *Lyve1^+/Cre^;R26R-tdTomato* embryos (A). tdTomato was found to be expressed not only in coronary blood and lymphatic vessels, but also in tissue resident macrophages (B). CD68^+^ cardiac resident macrophages appear to co-express CD206 and tdTomato, while they were found to be either LYVE1 positive (white arrows) or LYVE1 negative (hollow arrows) at E14.5 (C-F). Macrophages on the dorsal side of the heart (G). Similar to the ventral side of the heart, macrophages were CD68^+^;CD206^+^;tdTomato^+^and either LYVE1^+^ (white arrows) or LYVE1^−^ (H-L; hollow arrows). Quantification of the cardiac macrophage population found that 54 % were positive for LYVE1 and 44% were negative for LYVE1 (M). A small population of CD68^+^;CD206^−^;tdTomato^−^macrophages was found to contribute 2 % to the heart at E14.5. B-F magnified view of A box; H-L magnified view of G box. n = 3. Scale bars: 0.5 mm for A and G; 0.2 mm for B-F.

Taken together, *Lyve1* drives Cre activity in cardiac lymphatic vasculature during embryogenesis and additionally marks the sinus venosus, endocardium, and tissue-resident macrophages contributing to coronary vasculature and valve remodeling.

### Conditional deletion of *Prox1* in the *Lyve1^+^* lineage leads to impaired development and blood-filled cardiac lymphatics

Following detailed characterization of the *Lyve1^Cre^*driver, the *Prox1^Flox^* allele was employed to conditionally delete *Prox1* in the *Lyve1^+^* lineage and generate embryos with defective cardiac lymphatics. Previous reports described *Lyve1^+/Cre^;Prox1^+/Flox^*embryos as edematous, with most pups dying shortly after birth from peritoneal chylous ascites and ∼80% succumbing within three months^41,42^. In contrast, *Lyve1^+/Cre^;Prox1^+/Flox^* mice in our study exhibited no overt developmental defects and were viable and fertile as adults. These discrepancies likely reflect differences in *Prox1^Flox^* alleles—our line deletes the entire coding sequence and replaces it with GFP^43^, whereas the earlier studies deleted only the Homeodomain, leaving other functional domains intact^41,42^ — or genetic background effects, as survival in the prior studies required a pure NMRI background^41,42^.

To achieve complete *Prox1* loss, the *Lyve1^+/Cre^* driver was crossed with *Prox1^Flox/Flox^*mice, generating *Lyve1^+/Cre^;Prox1^Flox/Flox^* mutants. Genotyping confirmed Mendelian ratios: at E16.5, 62% controls (22/35) and 38% mutants (13/35); at E18.5, 74% controls (17/23) and 26% mutants (6/23). These data indicate that the *Lyve1^+/Cre^;Prox1^Flox/Flox^* genotype is not embryonic-lethal. However, no mutants were recovered at birth across four independent litters (0/20), demonstrating perinatal lethality (Supplementary Table 1-3).

Bright-field microscopy at E16.5 revealed subcutaneous oedema and blood-filled lymphatic vessels in the head, tail, and heart of *Lyve1^+/Cre^;Prox1^Flox/Flox^*embryos, phenotypes absent in littermate controls (Supplementary Figure 4A–L *vs* M–R). These findings indicate that lymphatic vessels are present in mutant hearts but fail to function properly, lacking separation from the blood vasculature. Mutant hearts also appeared paler than controls (Supplementary Figure 4C–F, I–L *vs* O–R), consistent with impaired lymphatic function. In contrast, no gross differences in overall body or heart size were detected between mutants and controls (Supplementary Figure 4A–L *vs* M–R).

To further define the vascular phenotype of *Lyve1^+/Cre^;Prox1^Flox/Flox^*embryos, immunostaining for EMCN and LYVE1 combined with light-sheet microscopy was performed at E16.5 (Figure 4A–L). In controls, cardiac lymphatics extended normally from base to apex along both dorsal and ventral surfaces of the heart (Figure 4B,E). In contrast, *Lyve1^+/Cre^;Prox1^Flox/Flox^* mutants lacked lymphatic vessels on the ventral aspect (Figure 4K) and displayed only sparse LYVE1^+^ structures on the dorsal side proximal to the sinus venosus (Figure 4H). Importantly, EMCN staining patterns were comparable between controls and mutants (Figure 4C–F *vs* I–L), indicating that coronary veins were unaffected. These findings demonstrate that conditional deletion of *Prox1* selectively impairs cardiac lymphatic development without altering coronary venous architecture.

**Figure 4.**
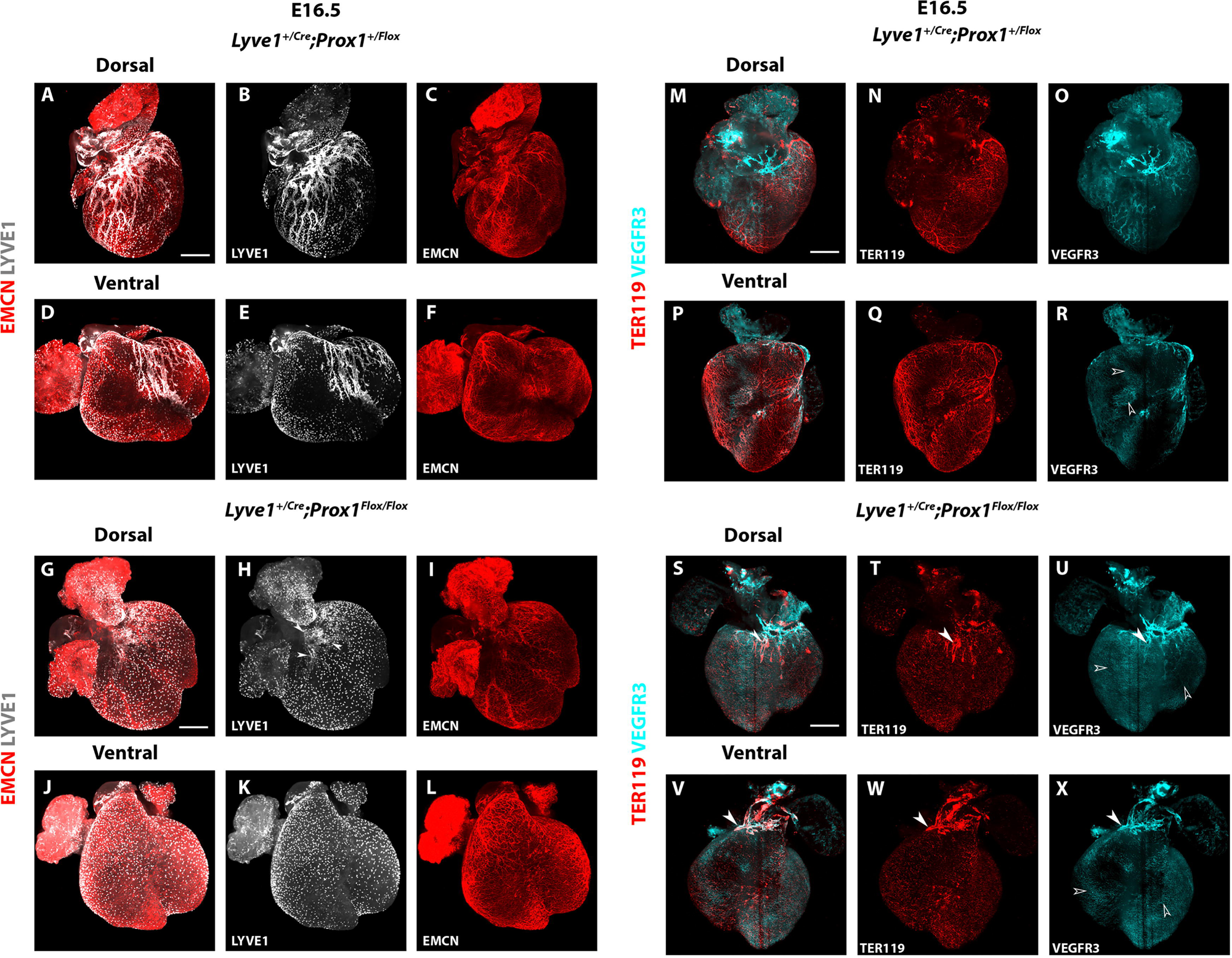
*Lyve1^+/Cre^;Prox1^Flox/Flox^*embryos have underdeveloped blood-filled cardiac lymphatics at E16.5. Dorsal side of a control *Lyve1^+/Cre^;Prox1^+/Flox^* heart stained against EMCN (venous and capillary endothelium) and LYVE1 (lymphatic endothelium and macrophages) (A-C). Lymphatic vessels extend from the SV towards the apex, while covering a large area of the ventricular surface. Also, individual LYVE1-expressing macrophages are visible (B). Blood vessels are covering the entire heart (C). Ventral side of the control heart shows extended and well developed lymphatic and blood vessels (D-F). In the dorsal side of a representative *Lyve1^+/Cre^;Prox1^Flox/Flox^* mutant heart, lymphatic vessels are underdeveloped and retained in the SV (arrowheads in panel H), while the blood vasculature appears grossly normal (G-I). In the ventral side of the heart, no lymphatics have formed, with LYVE1 signal being restricted to macrophages (J-L). The blood vasculature has no obvious defects (L). Dorsal aspect of a control *Lyve1^+/Cre^;Prox1^+/Flox^* heart stained with TER119 (erythrocytes) and VEGFR3 (lymphatics) (M-O). TER119 staining pattern represents erythrocyte-enriched blood vessels distributed throughout the heart alongside lymphatics (M-N). Lymphatics sprout from the SV and cover a large area of the ventricular surface (O). Ventral aspect of a control heart showing normal distribution of erythrocytes in blood vessels that do not mix with lymphatics (P-R). Note that at this timepoint VEGFR3 is still visible in blood capillaries (hollow arrowheads in panels R, U and X). On the dorsal side of a representative mutant *Lyve1^+/Cre^;Prox1^Flox/Flox^*heart, TER119 is clearly seen inside the few lymphatics located near the SV (white arrowheads) (S-U). Similarly, TER119 signal colocalized with VEGFR3 signal found in underdeveloped lymphatics in the outflow track (V-X; white arrowheads). n = 5 hearts per group. Scale bar: 0.5 mm.

To determine whether blood had infiltrated cardiac lymphatics in *Lyve1^+/Cre^;Prox1^Flox/Flox^* embryos, light-sheet microscopy was performed at E16.5 on hearts stained for VEGFR3 and TER119, an erythroid marker (Figure 4M–X). In controls, VEGFR3^+^ lymphatic vessels were well developed and extended across the dorsal and ventral heart surfaces (Figure 4O,R). TER119 signal was distributed throughout the myocardium (Figure 4N,Q) in close proximity to lymphatics, but never colocalized (Figure 4M,P), consistent with the parallel development of lymphatics and coronary veins. In contrast, *Lyve1^+/Cre^;Prox1^Flox/Flox^* mutants displayed severely underdeveloped lymphatics restricted to the cardiac base (Figure 4U,X). Strikingly, TER119 signal was enriched in VEGFR3+ regions (Figure 4S,V), confirming that mutant cardiac lymphatics were blood-filled at E16.5.

Surprisingly, the phenotype of *Lyve1^+/Cre^;Prox1^Flox/Flox^* embryos appeared less severe at E18.5 compared to E16.5 (Supplementary Figure 5). No visible edema or blood-filled lymphatics were detected, and the size and morphology of mutant embryos and their hearts were indistinguishable from littermate controls (Supplementary Figure 5A–F *vs* G–L). To assess whether cardiac lymphatics could recover or expand at this stage, light-sheet imaging was performed on E18.5 hearts stained for EMCN and LYVE1 (Figure 5A-L). As at E16.5, no cardiac lymphatics were observed in mutants, with LYVE1 signal restricted to cardiac macrophages (Figure 5H,K). In contrast, controls displayed normal base-to-apex expansion of cardiac lymphatics (Figure 5B,E). EMCN-labelled coronary veins showed no morphological differences between mutants and controls (Figure 5C,F *vs* I,L).

**Figure 5.**
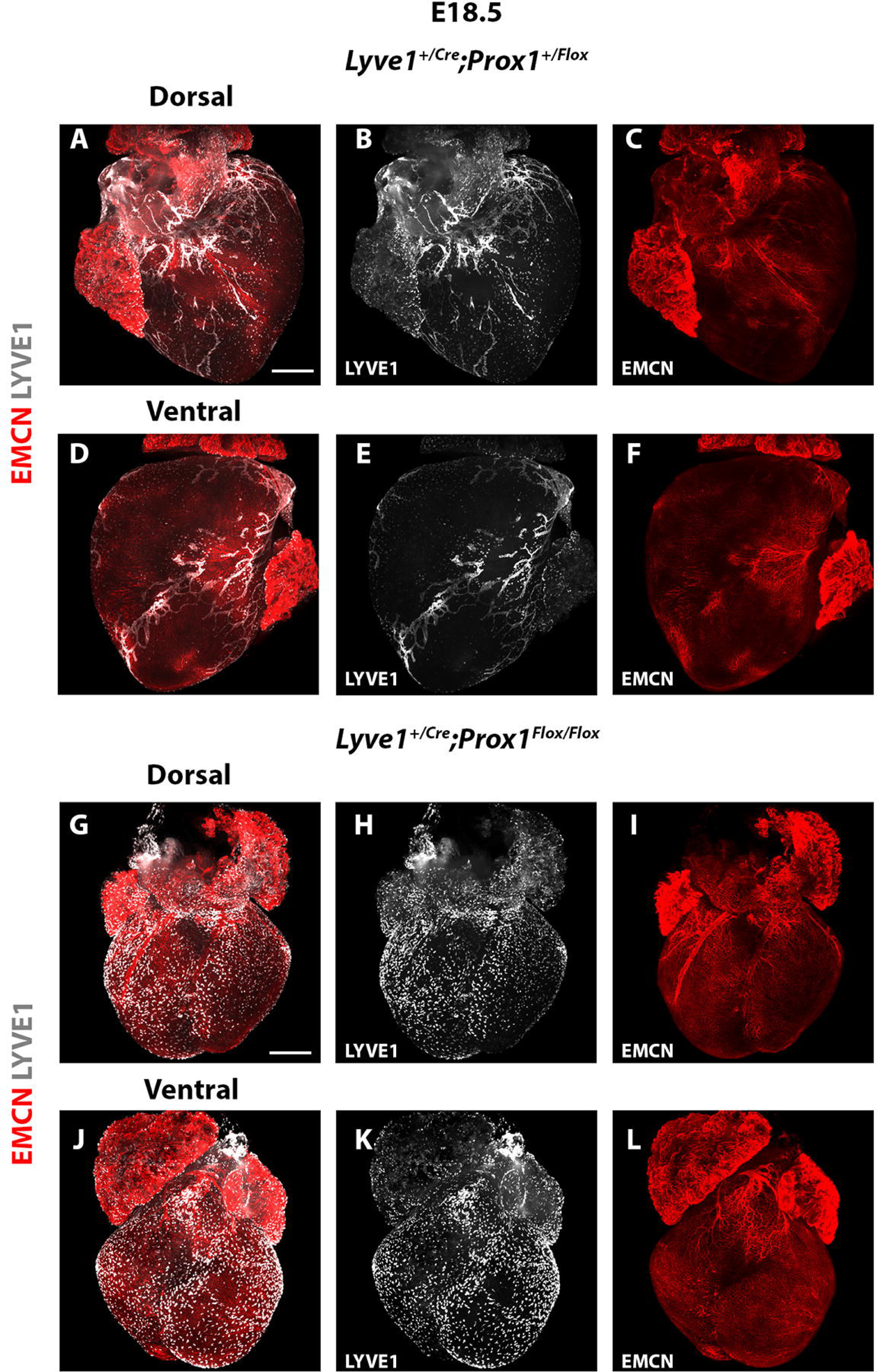
Cardiac lymphatics of *Lyve1^+/Cre^;Prox1^Flox/Flox^* embryos remain underdeveloped at E18.5. Dorsal side of a control *Lyve1^+/Cre^;Prox1^+/Flox^*heart (A-C). Lymphatics have developed through lymphangiogenesis and are covering most of the areas of the heart from SV to the apex (B). Blood vessels are covering the entire heart, with extended number of capillaries being present (C). Ventral side of the control heart shows normal lymphatic and blood vessels (D-F). On the dorsal side of a representative mutant *Lyve1^+/Cre^;Prox1^Flox/Flox^* heart no LYVE1^+^ lymphatics are visible, while the blood vasculature remains normal (G-I). Similarly, no lymphatics have formed on the ventral side of the heart, with the anti-LYVE1 antibody staining macrophages only (J-L). The blood vasculature has no obvious developmental defects (L). n = 6 hearts per group. Scale bar: 0.5 mm.

These findings demonstrate that the lymphatic phenotype observed at E16.5 is not rescued at later stages of fetal development. Instead, the absence of functional cardiac lymphatics persists and results in perinatal lethality, precluding the use of this mutant line for studies of neonatal cardiac regeneration.

High-resolution episcopic microscopy (HREM)^44,45^ was performed at E15.5 to assess cardiac morphology in *Lyve1^+/Cre^;Prox1^Flox/Flox^* embryos (Supplementary Figure 6A-O). All mutant hearts and their myocardial walls appeared smaller than controls (Supplementary Figure 6A *vs* E). Three of four mutants exhibited open tricuspid and mitral valve leaflets (Supplementary Figure 6B–D *vs* F-G; H *vs* I), consistent with valve malformation, while one mutant displayed a perimembranous ventricular septal defect (Supplementary Figure 6A *vs* E). The variability of these defects indicates incomplete penetrance of the phenotype in *Lyve1^+/Cre^;Prox1^Flox/Flox^* mice. Quantitative analysis of 3D HREM reconstructions confirmed significant size reductions in ventricles (*p* = 0.028), atria (*p* = 0.004), and tricuspid valves (p = 0.018) compared to controls (Supplementary Figure 6J–L). No statistically significant differences were detected in mitral, pulmonary, or aortic valves (Supplementary Figure 6M–O).

Altogether, these results demonstrate *Lyve1^Cre^* activity in the coronary vasculature, lymphatic progenitors, endocardium, and tissue-resident macrophages, and that *Prox1* is indispensable for cardiac lymphatic separation from blood vessels and valvulogenesis, with its loss resulting in blood-filled lymphatics and perinatal lethality.

## Discussion

This study establishes *Lyve1^Cre^* as a versatile but non-specific driver of cardiovascular lineage tracing, revealing activity across venous endothelium, endocardium, lymphatic vasculature, and resident macrophages. Previous reports have described *Lyve1^Cre^* activity in yolk sac hemogenic endothelium and subsets of endothelial cells^16^, but its relevance to the embryonic heart has not hitherto been comprehensively defined. Our findings highlight both the utility of this line in tracing multiple cardiovascular cell types during development and the associated challenges in interpreting loss-of-function phenotypes with the *Lyve1^Cre^*-driver, which may reflect contributions from multiple lineages rather than lymphatic endothelium alone. Our lineage-tracing experiments demonstrate that *Lyve1^Cre^* marks coronary vessels derived from both sinus venosus and endocardium, thereby implicating dual progenitor sources in coronary plexus formation. This dual contribution aligns with current understanding of coronary vessel ontogeny, where both venous and endocardial contributions are now well established^29–33^. The detection of *Lyve1^Cre^*-labelled macrophages in valves further expands its scope, linking endocardial-derived immune cells to valve remodeling. This observation reinforces emerging evidence that immune–endothelial interactions are integral to cardiac morphogenesis^46^, consistent with recent reports of macrophage involvement in valve maturation and homeostasis^35^.

The observation of heterogeneous Lyve1^+^ and Lyve1^−^ resident macrophages is heterogeneity, consistent with yolk sac–derived origins and suggests functional diversification within the macrophage pool that may influence tissue homeostasis and repair. Comparable heterogeneity has been reported in other tissue-resident macrophage populations, where lineage origin and transcriptional state dictate functional specialization^46,47^. Our findings therefore position *Lyve1^Cre^* as a useful, albeit broad, tool for dissecting macrophage contributions to cardiovascular development, while also cautioning that lineage tracing may capture overlapping populations with distinct roles.

In the lymphatic compartment, *Lyve1^Cre^* incompletely labelled *Prox1*^+^ precursors at early stages, with full labelling only achieved as lymphatics matured. This temporal heterogeneity implies additional sources of cardiac lymphatics, potentially including SHF derivatives, and is consistent with the now accepted view that cardiac lymphatics arise from multiple progenitor populations beyond venous endothelium^4,8–10^. Such complexity echoes recent lineage-tracing studies that have identified multiple progenitor contributions to lymphatic networks in other organs^48–50^. Conditional *Prox1* deletion in *Lyve1^+^* lineages revealed the indispensability of *Prox1* for lymphatic separation and valvulogenesis, with perinatal lethality underscoring its developmental importance. The collection of defects, notably blood-filled lymphatics, valve malformations, and septal abnormalities reflects the intersection of lymphatic and endocardial dysfunction, highlighting *Prox1* as a central regulator of cardiovascular morphogenesis.

Together, these findings significantly enhance our understanding of *Lyve1^Cre^* as a valuable lineage-tracing tool and highlight the importance of careful intrepertation of loss-of-function phenotypes associated with targeting Lyve1^+^ lineages. They establish *Prox1* as essential for lymphatic and valvular integrity, while revealing unexpected contributions of *Lyve1^+^* macrophages and endocardial derivatives to cardiac development. Future studies employing inducible Cre drivers, such as *Prox1-CreERT2*, will be critical to disentangle lineage-specific roles and to probe how lymphatic dysfunction shapes perinatal survival and neonatal cardiac regeneration^24^.

More broadly, our work places cardiac lymphatic biology within the wider framework of cardiovascular development, emphasizing the need to integrate immune, blood and lymphatic endothelial cell components in interpreting the morphogenesis of the heart and has likely implications for understanding mechanisms of cardiac repair and regeneration.

## Acknowledgments

We thank staff at the University of Oxford Biological Services Building for animal husbandry and the Micron Bioimaging facility for access to advanced confocal and light-sheet microscopy. This work was funded by the British Heart Foundation (BHF Chair Award to PRR: CH/11/1/28798; BHF Programme Grant to PRR: RG/18/33532; BHF Intermediate Basic Science Research Fellowship to JMV: FS/19/31/34158), MRC Unit funding (MC_UU_00008/2) and BBSRC funding (BB/X007278/1) to DGJ, and was supported by Wellcome Trust PhD Program in Chromosome and Developmental Biology to KK.

## Author Contributions

JMV and PRR conceived the original project, sourced funding, and supervised the project direction and data analyses. KK and KZ carried out experiments and analyzed the data. JKS and DS carried out HREM and assisted with data analysis. NS provided the *Notch1* gain-of-function mouse model. DGJ provided the *Lyve1^Cre^* mouse strain and provided advice and technical assistance. JMV wrote the initial draft of the manuscript. PRR and JMV contributed to editing of the final manuscript.

## Methods

### Mouse Strains

Genetically modified mouse lines used in the study were kept in a pure C57BL/6 genetic background. The following mouse strains were used: *Prox1^loxP/^*^+*43*^, *Lyve1^EGFP-hCre/^*^+*14*^, *R26R-tdTomato^51^*, *Gt(ROSA)26Sor^tm1(Notch1)Dam^*^36^. Breeding was carried out using only *Cre^+^* males for all Cre strains. Mice were cared for and housed by Oxford University Biomedical Services. Investigators were blinded to genotype groups. All animal experiments were carried out according to UK Home Office project licenses PPL PC013B246 and PDDE89C84 and were compliant with the UK Animals (Scientific Procedures) Act 1986.

### Timed matings

To generate embryos, female mice were paired with male studs and were checked for vaginal plugs each morning. The day the vaginal plug was observed was designated as embryonic day 0.5 (E0.5).

### Embryonic heart dissection

Embryos were harvested from pregnant females at the required embryonic stage. The female was euthanized by cervical dislocation, and the uterine horns were dissected from the abdominal cavity. Embryos were extracted from the uterine horns, placed in ice cold Phosphate-buffered saline (PBS) (Sigma) and the amniotic sac was removed. Hearts were dissected from embryos for subsequent experimental analysis.

### Cryo-sectioning

Whole embryo and embryonic hearts were fixed in 4 % PFA overnight at 4°C. Following fixation, samples were washed three times for 10 min in PBS and then transferred to 30 % sucrose and PBS overnight at 4°C. Then, the samples were equilibrated in a 1:1 solution of OCT and 30 % sucrose for 1 hour in 4°C. After equilibration the hearts were embedded in 100 % OCT and frozen at −80°C. 20-25 μm slices were cut using a cryostat and transferred onto Superfrost Plus slides (VMR). Slides were dried on a slide dryer for 15 min before being rinsed with PBS.

### Immunostaining

Sections underwent permeabilization with 0.5 % Triton X-100 (Sigma, UK) for 10 min, followed by two rinses in PBS for 5min. Then, sections were blocked in blocking solution, composed of 10 % serum, 4 % bovine serum albumin (BSA) and 0.2 % Triton X-100, for 1 hour. The blocking serum was from the same species in which the secondary antibodies were made. Blocking was followed by a 4°C overnight incubation in primary antibody, which was diluted in blocking solution. A list of primary antibodies and the dilutions used are included in Table 1. For each primary antibody used, one section was incubated without primary antibody as a secondary antibody alone control. Following primary antibody incubation, sections were washed several times with 0.1 % Triton X-100 and incubated with Alexa Fluor-conjugated secondary antibodies for 1 hour at room temperature in the dark. All secondary antibodies were diluted in PBS; a list of the secondary antibodies used is included in Table 2. Following incubation in secondary antibody, the slides were washed several times in 0.1 % Triton X-100, with DAPI included in the final 15 min wash. A small amount of 50:50 Glycerol/PBS was then added to the slides and a 22×50 mm coverslip (Fisherbrand) was placed on top and sealed with nail varnish. Immunofluorescent staining was imaged using a Zeiss LSM780, Zeiss LSM880, Zeiss LSM980 or Leica confocal microscope. Z-stack and tiling functions were used when required. Images were processed using ImageJ software^52,53^.

**Table 1.** List of primary antibodies.

| **Primary antibody** | **Company/Catalogue number** | **Tissue expression/marker** | **Dilution** | **Species raised in** |
| --- | --- | --- | --- | --- |
| VEGFR3 | R&D systems<br>#AF743 | Lymphatic endothelium | 1:50 | Goat |
| PROX1 | R&D Systems<br>#AF2727 | Lymphatic endothelium and cardiomyocytes | 1:50 | Goat |
| LYVE1 | Angiobio<br>#11-034 | Lymphatic endothelium, tissue-resident macrophages, and endocardium | 1:400 | Rabbit |
| PODOPLANIN<br>(PDPN) | Fitzgerald #10R-P155A | Lymphatic<br>endothelium and<br>epicardium | 1:200 | Hamster |
| PECAM1 | BD Pharmingen<br>#553370 | Endothelium | 1:200 | Rat |
| ENDOMUCIN<br>(EMCN) | Santa Cruz<br>Biotechnology #SC-53941 | Venous and<br>capillaries<br>endothelium | 1:200 | Rat |
| TER119 | BD Biosciences<br>#550565 | Erythrocytes | 1:200 | Rat |
| CD68 | Bio-Rad #MCA1957 | Macrophages | 1:400 | Rat |
| CD206 | R&D Systems<br>#AF2535 | Tissue-resident<br>macrophages | 1:200 | Goat |

**Table 2.**
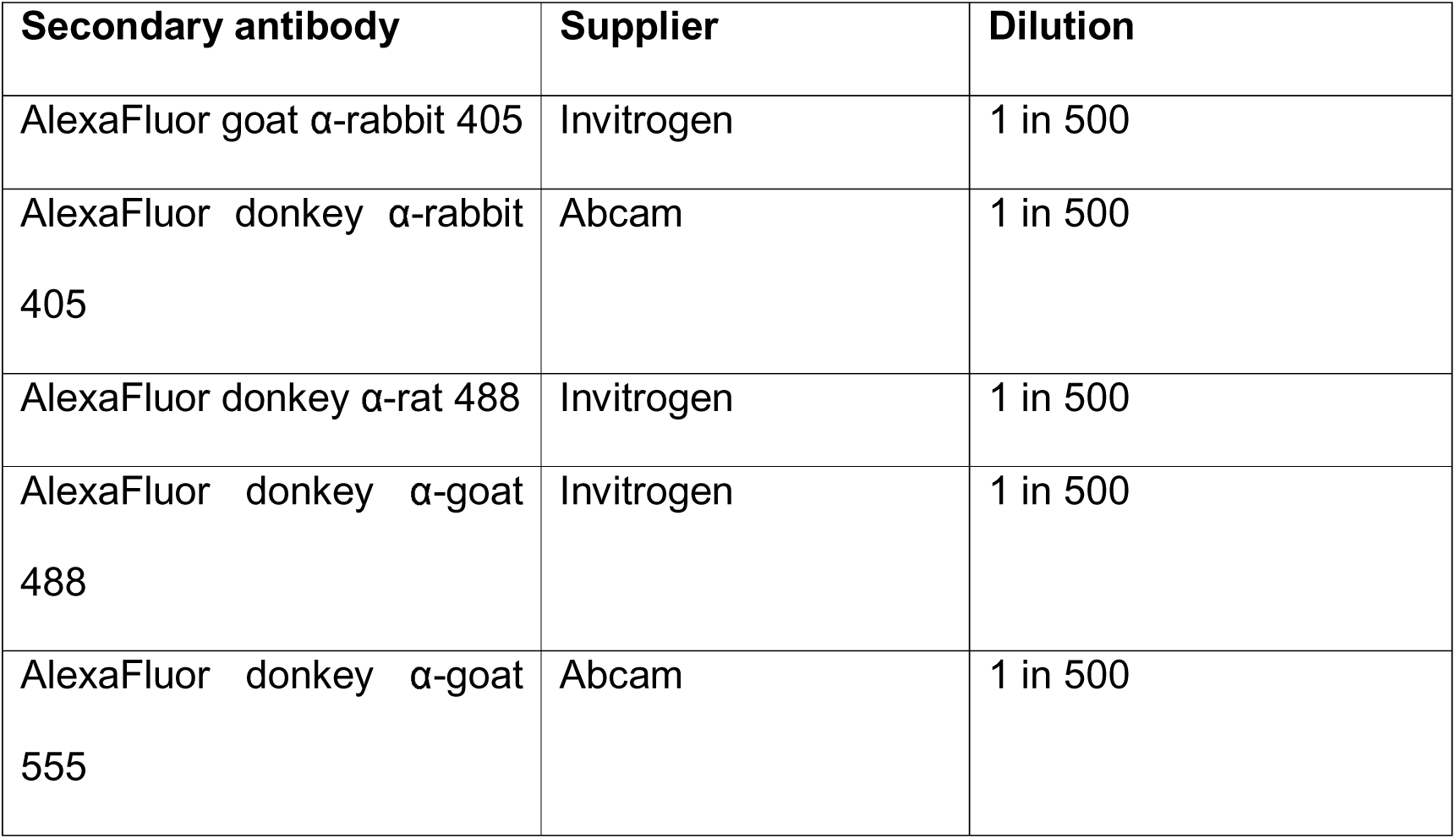
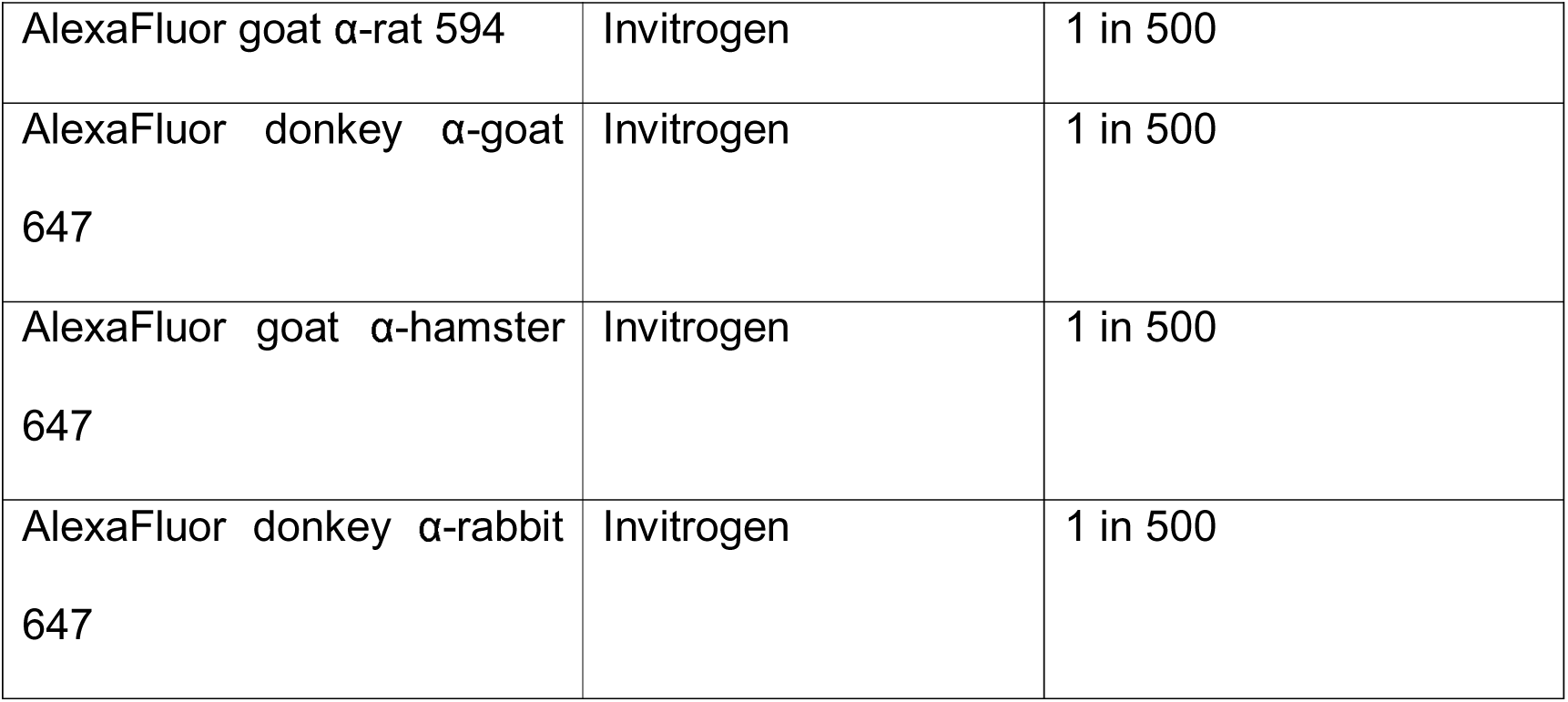
List of secondary antibodiess.

| Secondary antibody | Supplier | Dilution |
| --- | --- | --- |
| AlexaFluor goat α-rabbit 405 | Invitrogen | 1 in 500 |
| AlexaFluor donkey α-rabbit<br>405 | Abcam | 1 in 500 |
| AlexaFluor donkey α-rat 488 | Invitrogen | 1 in 500 |
| AlexaFluor donkey α-goat<br>488 | Invitrogen | 1 in 500 |
| AlexaFluor donkey α-goat<br>555 | Abcam | 1 in 500 |
| AlexaFluor goat α-rat 594 | Invitrogen | 1 in 500 |
| AlexaFluor donkey α-goat 647 | Invitrogen | 1 in 500 |
| AlexaFluor goat α-hamster 647 | Invitrogen | 1 in 500 |
| AlexaFluor donkey α-rabbit 647 | Invitrogen | 1 in 500 |

### Immunostaining of embryonic hearts

Whole embryonic hearts were permeabilized for 1 hour in 4 % Triton X-100 and subsequently blocked in blocking solution (2 % BSA, 10 % serum, 0.2 % Triton X-100 in PBS) overnight at 4°C. The blocking serum was from the same species in which the secondary antibodies were made. Samples were incubated with primary antibodies (**Table** 2) diluted in blocking solution for 48 hours at 4°C, then washed ten times for 30 min in PBS. Then, samples were incubated with secondary antibodies (Table 3) diluted in PBS overnight at 4°C in the dark. The hearts were washed five times for 15 min in PBS, with DAPI staining included in the last 30 min wash. Immunofluorescent staining was imaged using Zeiss LSM780, Zeiss LSM880, Zeiss LSM980 or Leica confocal microscope, or a Zeiss Z.1 light-sheet microscope. Z-stack and tiling functions were employed to obtain maximum intensity Z-projections of whole hearts. Images were processed using Imaris, Arivis Vision4D and ImageJ software.

For confocal imaging, samples were mounted on glass slides (VWR) covered in 3-4 layers of electrical tape with windows cut into the tape to create wells. The samples were covered with 50:50 Glycerol/PBS and a coverslip (Fisherbrand) was placed on top of the slides.

For light-sheet imaging, embryonic hearts were mounted using 1 mL syringes. A gel solution of 1.5 % low melting point agarose in TBE was prepared, and the sample was positioned inside the gel. The tip of the syringe was cut off to create an even cylinder and the gel solution with the sample was pumped in using the plunger. After the gel polymerized, the sample was pushed out and was ready for imaging.

### High-Resolution Episcopic Microscopy (HREM)

E15.5 embryos were fixed overnight in 4 % paraformaldehyde at 4°C and were dissected for heart-lung complexes. The heart-lung complexes were fixed in 4 % PFA and dehydrated by performing serial washes, each for 1 hours, in the following dilutions of methanol: 10 %, 20 %, 30 %, 40 %, 50 %, 60 %, 70 %, 80 %, 90 %, 95 % and 100 %. The samples were incubated in 50:50 mixtures of methanol/JB4 resin (Polysciences, 00226-1, GMBH, Germany) overnight. The samples were incubated in JB4 resin for 1 hour and transferred to fresh resin for 3 days’ incubation. The samples were embedded individually in JB4 according to manufacturer’s instructions and were cut at 3 μm sections on an optical HREM (OHREM) microscope (Indigo scientific) with images taken using a Jenoptik Gryphax camera. Image stacks were processed into cubic data and reduced to 50% for 3D modelling using the Amira software package 2019.4 (Thermo Fisher Scientific). Greyscale images were imported into MD DICOM viewer version 9.0.2 (Pixmeo), or Horos 3.3.6 (https://horosproject.org) in a 50% stack, inverted to black-white, and rendered into 3D.

### Statistical analysis

All statistical analyses were performed using GraphPad Prism 8 software. Comparisons between two groups were made using an unpaired two-tailed T test, this included an F test to confirm the two groups had equal variances. In all cases a *p-value* of less than or equal to 0.05 was deemed significant.

