## Supplementary Figures 1-6; Supplementary Tables 1-3 for "Distinct roles of the *Lyve1* lineage in heart development"

### Supplementary information

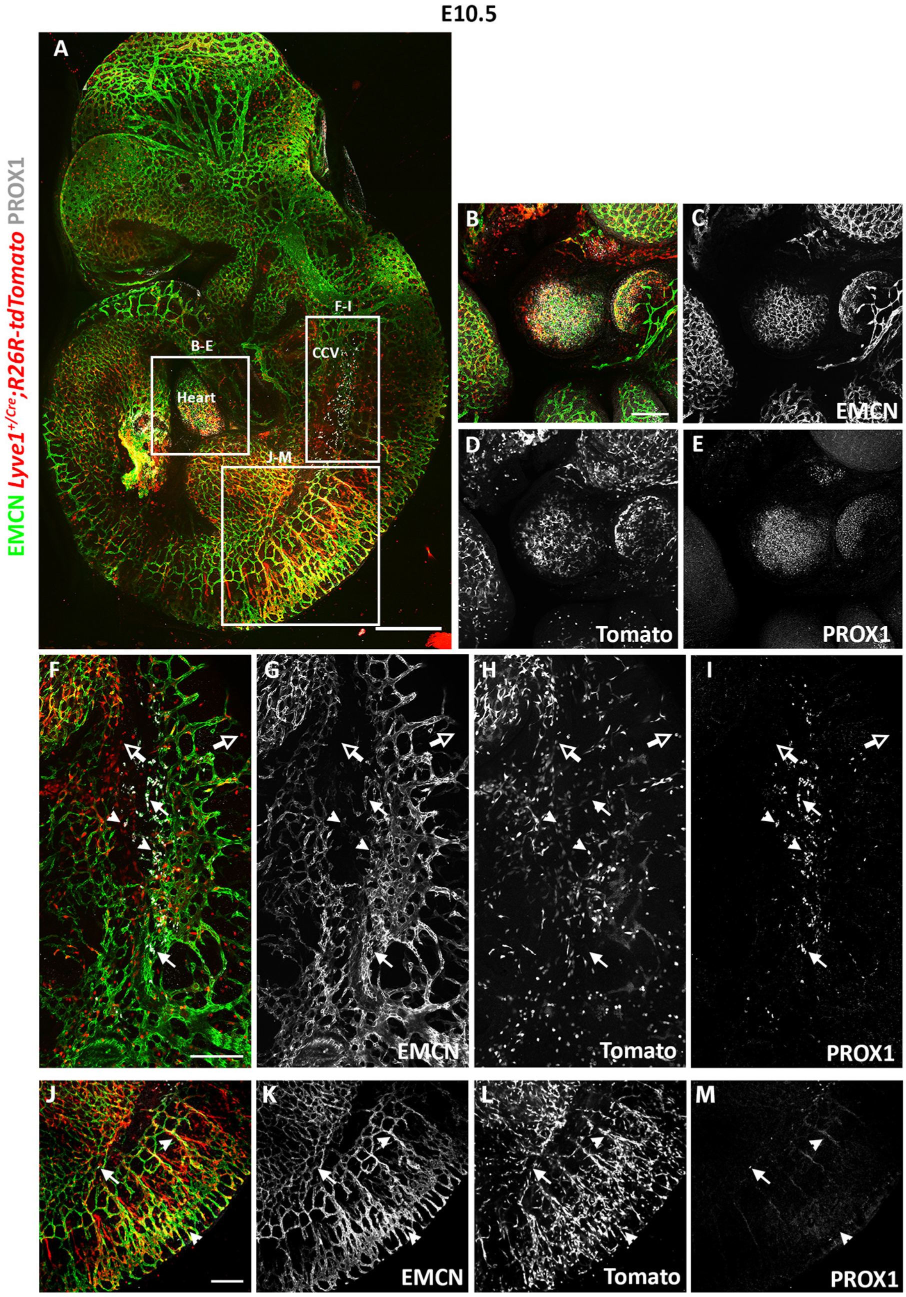


**Supplementary Figure 1. *Lyve1$^{Cre}$* is activated in intersomitic vessels, heart capillaries and partially in LEC precursors of the common cardinal vein.**

Representative E10.5 *Lyve1$^{+/Cre}$;R26R-tdTomato* embryo stained with anti-EMCN and anti-PROX1 antibodies (A). High magnification shows EMCN, PROX1 and tdTomato signal in the embryonic heart (B-E). High magnification of the CV reveals tdTomato signal in EMCN$^{+}$ vessels, while individual PROX1$^{+}$ precursor LECs are either EMCN$^{+}$;tdTomato$^{+}$ (arrowheads) or EMCN$^{+}$;tdTomato$^{-}$ (white arrows). Macrophages are also visible (hollow arrows) (F-I). High magnification of the ISVs displayed colocalization of EMCN and tdTomato signal in the vessels (arrowheads), as well as $PROX1^{+}$;$EMCN^{+}$;$tdTomato^{-}$ precursor LECs (white arrows) (J-M). B-E, F-I and J-K magnified views of A boxes. n = 3 embryos. Scale bars: 0.5 mm for A; 0.2 mm for B-M.

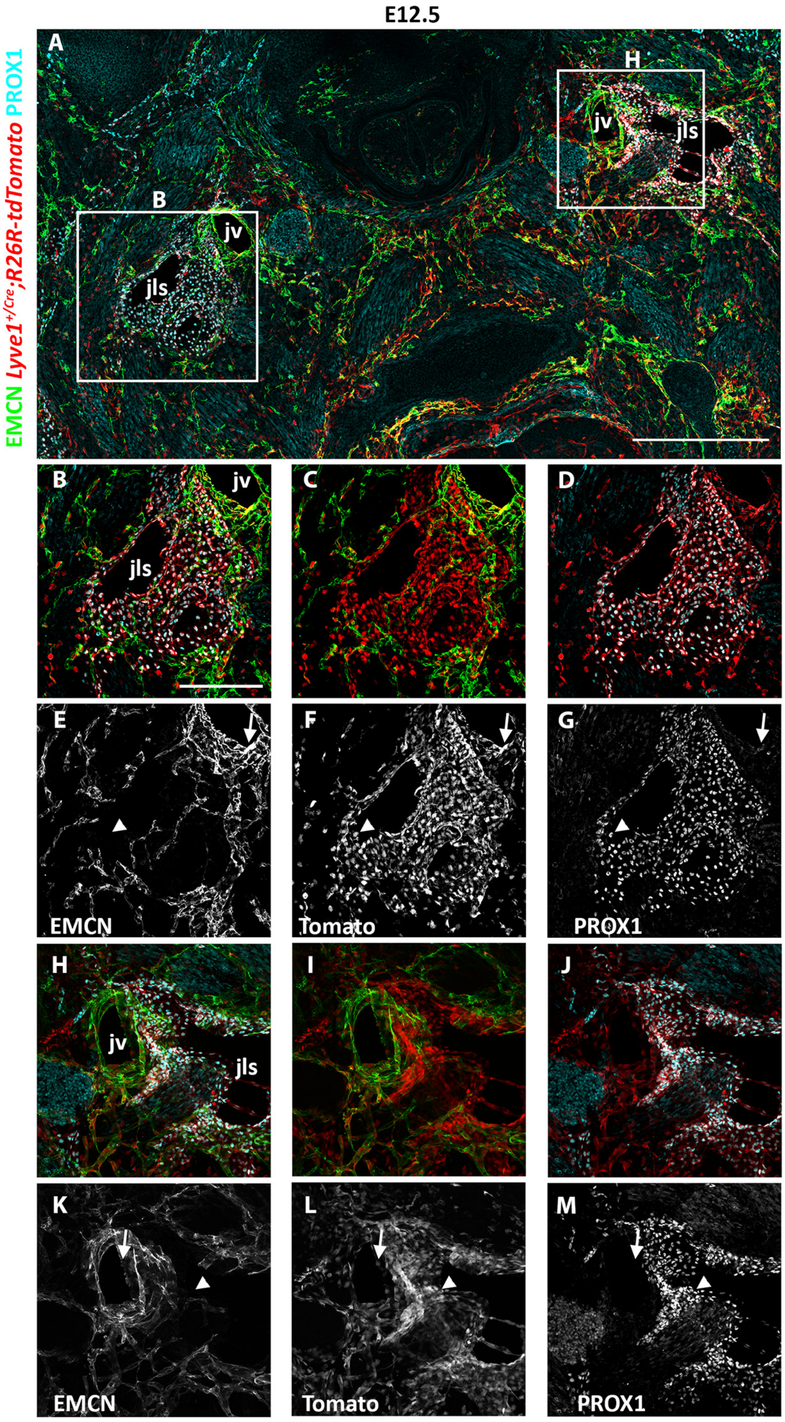


**Supplementary Figure 2. *Lyve1$^{Cre}$* is activated in both the jugular vein and jugular lymph sacs at E12.5.**

Transverse sections of *Lyve1$^{+/Cre}$;R26R-tdTomato* embryos and immunostaining with antibodies against EMCN and PROX1 at E12.5 (A). High magnification of the JLSs and JVs shows high levels of tdTomato expression in both EMCN$^-$;PROX1$^+$ (JLSs; arrowheads) and EMCN$^+$;PROX1$^-$ (JVs; white arrows) (B-M). B-G and H-M magnified views of A boxes. n = 3 embryos. Scale bars: 0.5 mm for A; 0.2 mm for B-M.

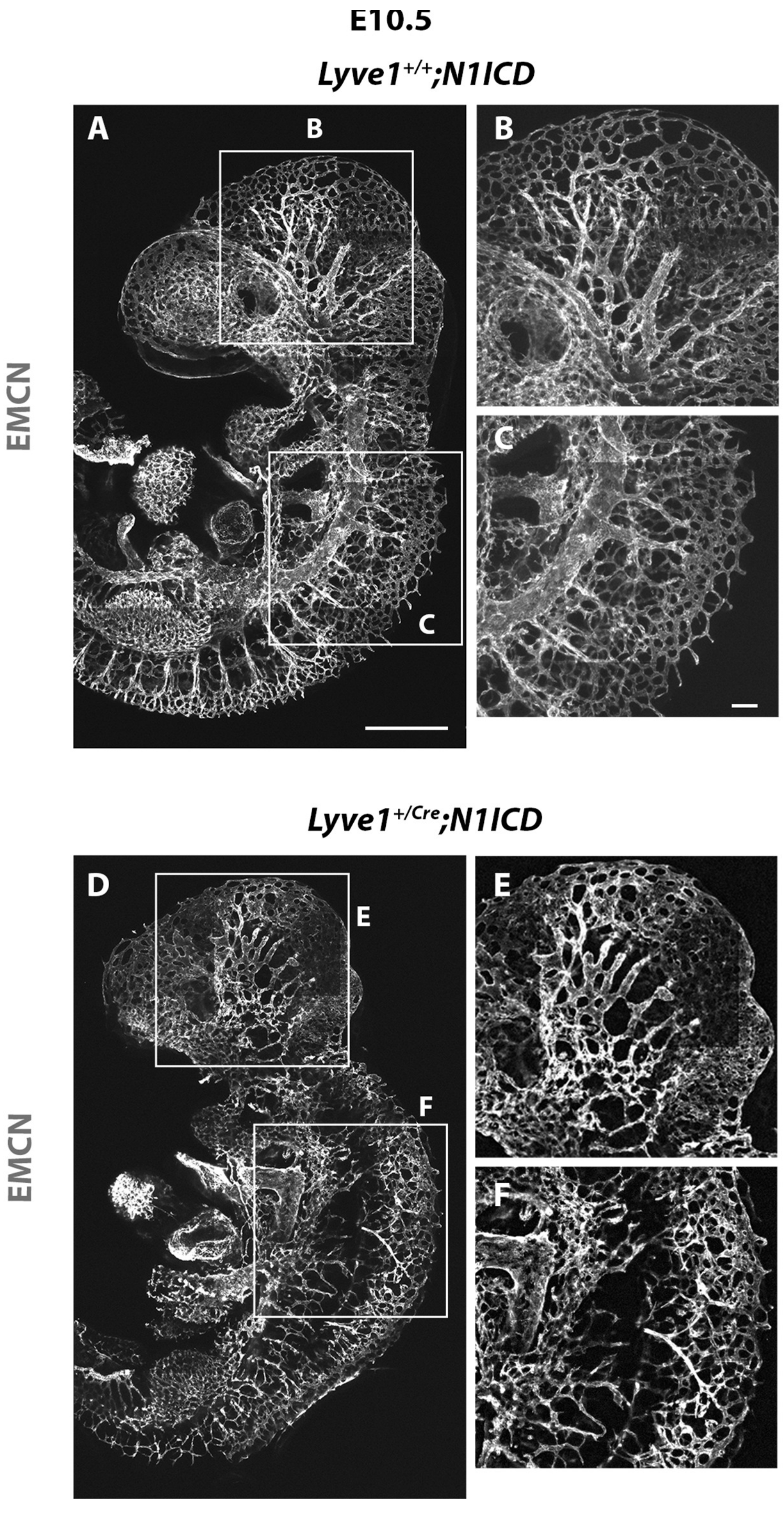


**Supplementary Figure 3. *Notch1* gain-of-function in *Lyve1*$^{+}$ lineage leads to vascular defects in the developing embryo.**

Confocal imaging of control *Lyve1*$^{+/+}$*;N1ICD* and mutant *Lyve1*$^{+/Cre}$*;N1ICD* embryos stained against EMCN at E10.5 (A-F). The blood vasculature appeared morphologically normal with branches and sprouts coming from larger vessels to thinner capillaries in the control embryos (A). Details of the well developing blood vasculature can be observed with high magnification in the ISVs and head of control embryos (B-C). Major defects were detected in the development of the blood vasculature in mutant embryos, particularly in the head and ISVs (D). In the head, the blood vessels had lower density, were thicker and showed reduced level of remodelling of the vascular plexus, compared to the control (E). The ISVs were disorganized and often failed to connect to the CV (F). B-C magnified views of A boxes; E-F magnified views of D boxes. n = 5. Scale bars: 0.5 mm for A and D; 0.2 mm for B-C and E-F.

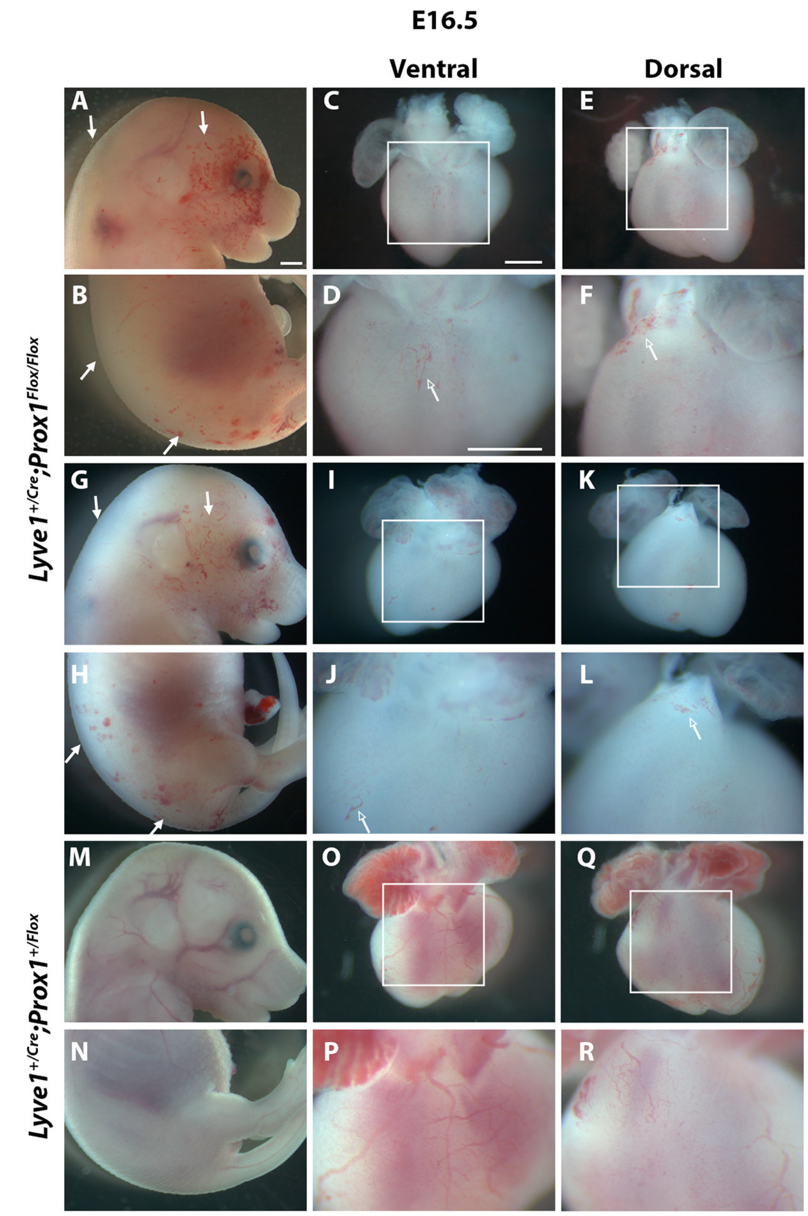


**Supplementary Figure 4. *Lyve1$^{+/Cre}$;Prox1$^{Flox/Flox}$* embryos display gross developmental defects at E16.5.**

Two representative mutant embryos and corresponding hearts (A-L). Both embryos have severe oedema and blood-filled lymphatic vessels in the head, back and tail (white arrows) (A-B and G-H). High magnification clearly shows that the heart of mutant embryos has blood-filled lymphatics near the base on the ventral and dorsal side (hollow arrows) (C-F and I-L). Control embryos do not have oedema, nor blood-filled lymphatics in the body or the heart (M-R). D and F magnified views of C and E boxes; J and L magnified views of I and K boxes; P and R magnified views of O and Q boxes. n = 22 littermate control; n = 13 mutants. Scale bars: 0.5 mm.

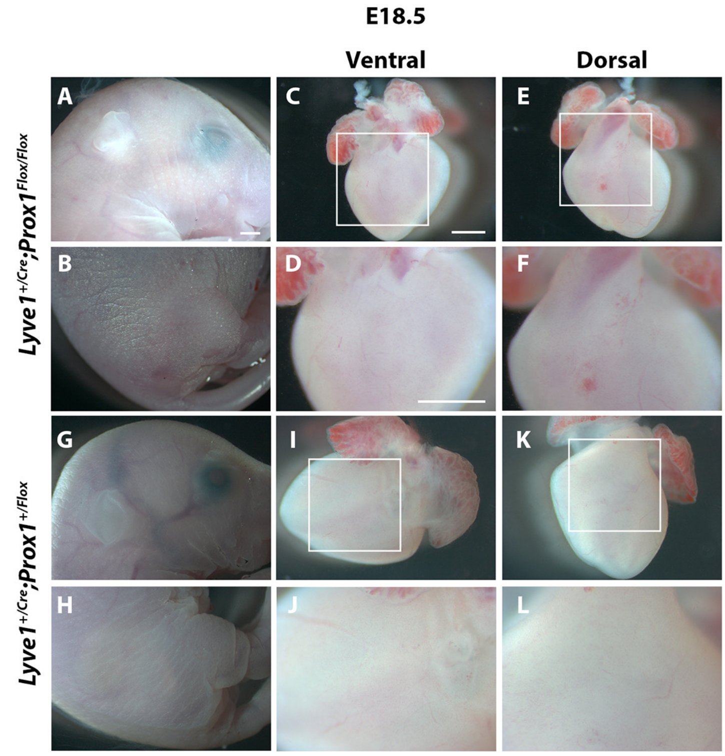


**Supplementary Figure 5.** ***Lyve1$^{+/Cre}$;Prox1$^{Flox/Flox}$*** **embryos display grossly normal development at E18.5.**

Representative mutant embryo and its heart (A-F). At this timepoint, mutant embryos do not exhibit oedema or blood-filled lymphatics (A-B). No obvious cardiac vessel defects are observed in mutant embryos at E18.5 (C-F). Control embryos do not exhibit oedema, nor blood-filled lymphatics in the body or heart (G-L). D and F magnified views of C and E boxes; J and L magnified views of I and K boxes. n = 17 littermate controls; n = 6 mutants. Scale bars: 0.5 mm.

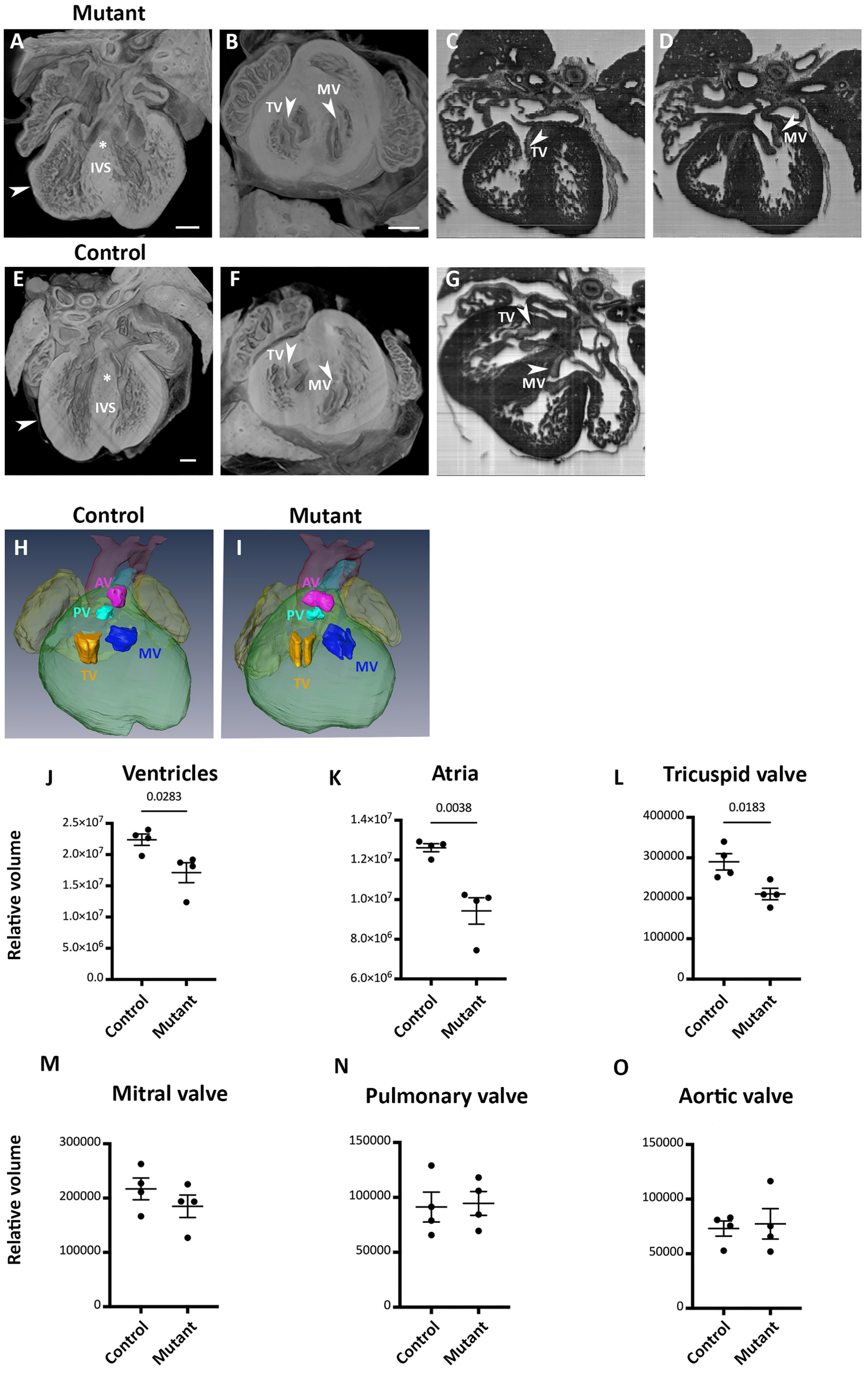
Mutant
A
*
IVS
B
TV
MV
C
TV
D
MV
Control
E
*
IVS
F
TV
MV
G
TV
MV
Control
Mutant
H
AV
PV
TV
MV
I
AV
PV
TV
MV
J
Ventricles
0.0283
Relative volume
2.5×10^7
2.0×10^7
1.5×10^7
1.0×10^7
5.0×10^6
0.0
Control
Mutant
K
Atria
0.0038
1.4×10^7
1.2×10^7
1.0×10^7
8.0×10^6
6.0×10^6
Control
Mutant
L
Tricuspid valve
0.0183
400000
300000
200000
100000
0
Control
Mutant
M
Mitral valve
Relative volume
300000
200000
100000
0
Control
Mutant
N
Pulmonary valve
150000
100000
50000
0
Control
Mutant
O
Aortic valve
150000
100000
50000
0
Control
Mutant

**Supplementary Figure 6. 3D HREM modelling of *Lyve1$^{+/Cre}$;Prox1$^{Flox/Flox}$* hearts shows developmental defects.**

One representative mutant heart showing developmental defects (A-D). The interventricular septum (IVS) is membranous (asterisk), and the myocardial wall is thin (white arrowhead) (A). The mitral (MV) and tricuspid (TV) valve leaflets are open only in mutant hearts (B-D). A representative control heart, displaying normal IVS, thick myocardial wall, and well closed MV and TV leaflets (E-F). Representative 3D rendering of a control and mutant heart. The leaflets of the TV are closed in the control compared to the mutant (H-I). Quantification of the 3D rendering showing significant smaller ventricles, atria, and tricuspid valve in mutant hearts compared to controls (J-O). Pulmonary valve (PV), aortic valve (AV). n = 4. Significant differences were calculated using an unpaired, 2-tailed Student's t test. Scale bar: 0.5 mm.

**Supplementary Table 1. Embryo genotypes from *Lyve1*$^{+/Cre}$*;Prox1*$^{+/Flox}$ × *Lyve1*$^{+/+}$*;Prox1*$^{Flox/Flox}$ crosses at E16.5.**

| **E16.5** | **Control** | **Mutant** |
|---|---|---|
| Litter #1 | 9 | 6 |
| Litter #2 | 5 | 3 |
| Litter #3 | 2 | 2 |
| Litter #4 | 6 | 2 |
| **Observed** | 22 | 13 |
| **Expected** | 26.25 | 8.75 |

**Supplementary Table 2.** Embryo genotypes from *Lyve1*$^{+/Cre}$;*Prox1*$^{+/Flox}$ × *Lyve1*$^{+/+}$;*Prox1*$^{Flox/Flox}$ crosses at E18.5.

| **E18.5** | **Control** | **Mutant** |
|---|---|---|
| Litter #1 | 5 | 4 |
| Litter #2 | 6 | 1 |
| Litter #3 | 6 | 1 |
| **Observed** | 17 | 6 |
| **Expected** | 17.25 | 5.75 |

**Supplementary Table 3.** Neonatal genotypes from *Lyve1*$^{+/Cre}$;*Prox1*$^{+/Flox}$ × *Lyve1*$^{+/+}$;*Prox1*$^{Flox/Flox}$ crosses.

| **Born** | **Control** | **Mutant** |
|---|---|---|
| Litter #1 | 5 | 0 |
| Litter #2 | 5 | 0 |
| Litter #3 | 4 | 0 |
| Litter #4 | 6 | 0 |
| **Observed** | 20 | 0 |
| **Expected** | 15 | 5 |
